# An angiosperm *NLR* atlas reveals that *NLR* gene reduction is associated with ecological specialization and signal transduction component deletion

**DOI:** 10.1101/2021.02.10.430603

**Authors:** Yang Liu, Zhen Zeng, Qian Li, Xing-Mei Jiang, Zhen Jiang, Ji-Hong Tang, Yan-Mei Zhang, Dijun Chen, Qiang Wang, Jian-Qun Chen, Zhu-Qing Shao

**Affiliations:** State Key Laboratory of Pharmaceutical Biotechnology, School of Life Sciences, Nanjing University, Nanjing, 210023, China; Jiangsu Key Laboratory for the Research and Utilization of Plant Resources, Institute of Botany, Jiangsu Province and Chinese Academy of Sciences, Nanjing, 210014, China

**Keywords:** *NLR* genes, plant disease resistance, *EDS1* gene family, gene family evolution, angiosperms

## Abstract

Nucleotide-binding site-leucine-rich repeat receptor (*NLR*) genes comprise the largest family of plant disease resistance genes. *NLR* genes are phylogenetically divided into the *TNL, CNL,* and *RNL* subclasses. *NLR* copy numbers and subclass composition vary tremendously across angiosperm genomes. However, the evolutionary associations between genomic *NLR* content and plant lifestyle, or between *NLR* content and signal transduction components, are poorly characterized due to limited genome availability. Here, we established an <u>an</u>giosperm *<u>N</u>LR* <u>a</u>tlas (ANNA, http://compbio.nju.edu.cn/app/ANNA/), which includes *NLR* genes from over 300 angiosperm genomes. Using ANNA, we revealed that *NLR* copy numbers differ up to 66-fold among closely related species due to rapid gene loss and gain. Interestingly, *NLR* contraction was associated with adaptations to aquatic, parasitic, and carnivorous lifestyles. The convergent *NLR* reduction in aquatic plants resembles the long-term evolutionary silence of *NLR* genes in green algae before the colonization of land. A co-evolutionary pattern between *NLR* subclasses and plant immune-pathway components was also identified, suggesting that immune pathway deficiencies may drive *TNL* loss. Finally, we recovered a conserved *TNL* lineage that may function independently of the RNL pathway. Our findings provide new insights into the evolution of *NLR* genes in the context of plant lifestyles and genome content variation.

## Introduction

Plants rely on a two-layered immune system to defend against various pathogens: the first immune-system layer, termed pathogen-associated molecular pattern (PAMP)-triggered immunity (PTI), recognizes PAMPs using cell surface-localized receptors; the second immune-system layer, known as effector-triggered immunity (ETI), detects effectors released by pathogens using intracellular disease resistance genes (*R* genes) (Wang et al., 2020). Several different types of *R* genes have been identified over the past twenty years (Kourelis and van der Hoorn, 2018). The largest *R* gene family is comprised of genes encoding nucleotide-binding site (NBS) and leucine-rich repeat (LRR) domain receptors, which are known as *NLR* or *NBS-LRR* genes (Kourelis and van der Hoorn, 2018). The plant-specific *NLR* gene family originated and diverged in the common ancestor of green plants (Shao et al., 2019). Three *NLR* gene subclasses, *TIR-NBS-LRR* (*TNL*), *CC-NBS-LRR* (*CNL*), and *RPW8-NBS-LRR* (*RNL*), have been characterized based on the N-terminal domains of the encoded NLR proteins: the Toll/Interleukin-1 receptor (TIR) domain, the coiled-coil (CC) domain, and the resistance to powdery mildew8 (RPW8) domain, respectively (Pan et al., 2000; Parker et al., 1997; Shao et al., 2014). Most CNL and TNL proteins function as pathogen detectors, either directly interacting with pathogen effectors or monitoring the state alteration of host proteins targeted by the effectors (Kourelis and van der Hoorn, 2018). The proteins encoded by the *RNL* genes are “helper” NLRs, which are involved in the downstream signal transduction of CNL and TNL proteins (Kourelis and van der Hoorn, 2018; Wang et al., 2020).

Over much of evolutionary time, *NLR* genes have been silent and have indeed been frequently lost in the green algae (Shao et al., 2019). However, after land colonization, the *NLR* genes expanded rapidly in green-plant genomes: angiosperm genomes often possess dozens to hundreds of *NLR* genes, most of which belong to the *TNL* and *CNL* subfamilies (Shao et al., 2016). Numbers of *NLR* genes may vary widely among genomes within the same family: two- to six-fold differences in gene number have frequently been reported (Luo et al., 2012; Qian et al., 2017; Shao et al., 2014; Tirnaz et al., 2020; Zhang et al., 2016). Indeed, a recent study revealed interspecies differences in *NLR* gene number as high as 20-fold among species in the Orchidaceae (Xue et al., 2020). This suggests that dramatic *NLR* gain or loss might occur rapidly after speciation. Pathogenic selection pressure may be the primary driving force underlying these *NLR* gene expansions. However, possible associations between *NLR* gene number and adaptations to specific lifestyles remain largely unexplored, primarily because few angiosperm genomes have been sequenced.

The absence of *TNL* genes from monocot genomes and several dicot genomes is another long-standing puzzle in *NLR* gene evolution (Bai et al., 2002; Collier et al., 2011). Several recent studies investigated the involvement of proteins from the two RNL lineages (*ADR1* and *NRG1*) in TNL-triggered immunity. These studies revealed that nearly all tested TNL proteins were functionally dependent on RNL proteins, predominantly from the NRG1 lineage, to confer full resistance against pathogens (Castel et al., 2019; Qi et al., 2018; Saile et al., 2020; Wu et al., 2019). Moreover, a pattern of co-absence was observed for *TNL* and *NRG1* genes in one monocot and two dicot species (Collier et al., 2011). The functional and evolutionary relationships among *TNL* and *RNL* genes provide intriguing clues concerning the mechanism of *TNL* loss.

In addition to RNL proteins, other key signaling elements from the ENHANCED DISEASE SUSCEPTIBILITY (EDS1) family are required for *TNL-*triggered disease resistance. The EDS1 family includes EDS1, PHYTOALEXIN DEFICIENT 4 (PAD4), and SENESCENCE ASSOCIATED GENE 101 (SAG101); in *Arabidopsis thaliana*, EDS1 forms heterodimers with SAG101 and with PAD4 (Lapin et al., 2020). When pathogens invade and TNL proteins are activated, the resulting signaling involves the enzymatic catalysis of NAD^+^, which is transmitted downward to EDS1 via an unknown mechanism. EDS1 then further transmits the signal to downstream RNL proteins to induce local cell necrosis and transcriptional responses (Lapin et al., 2019; Wan et al., 2019).

Interestingly, several plant species have been shown to lack both *SAG101* and *TNL* (Lapin et al., 2019). The concomitant loss of the *TNL* gene and its signal transduction partners suggests that genomic context alteration may lead to variations in *NLR* gene copy number. However, this pattern of *TNL* and *NRG1/SAG101* co-absence has been observed in only a few angiosperm genomes, corresponding to three independent losses (Collier et al., 2011; Lapin et al., 2019; Shao et al., 2016). It remains unclear how many other angiosperm genomes lack *TNL* genes, and whether these losses are related to similar genome content alterations.

Due to recent advances in sequencing technologies, hundreds of genomes, representing many of the major angiosperm clades, have been sequenced. This provides a unique opportunity to explore *NLR*-associated questions using a large dataset. Thus, in this study, we establish an angiosperm *NLR* atlas (ANNA) by identifying and incorporating over 90,000 *NLR* genes from more than 300 angiosperm genomes. Using this huge dataset, we determined that plants that adapted to aquatic, parasitic, and carnivorous lifestyles tended to lose *NLR* genes. In addition, numbers of *TNL* and *RNL* genes were highly correlated within a genome, and *TNL* loss tended to coincide with the loss of the *SAG101* gene, as well as genes in the *RNL-NRG1* lineage. This suggested that *NRG1/SAG101* pathway deficiency may potentially drive *TNL* loss. These findings provide new insights into genome context variation and the adaptive evolution of disease resistance genes to plant lifestyles.

## Results

### Construction of the ANNA, including over 300 genomes

We identified 91,291 *NLR* genes in 305 publicly available angiosperm genomes, including four basal angiosperms, 65 monocots, 230 dicots, five magnoliids, and one Ceratophyllales (Table S1, S2). These species were distributed across the major angiosperm clades, representing 85 families in 37 of the 64 angiosperm orders (Figure 1A). The 65 monocot genomes represented nine families in six orders, providing a large and diverse group of taxa with which to test the hypothesis of ancestral loss of *TNL* genes in this clade. The 230 dicot taxa comprised seven species representing two basal lineages (Proteales and Ranunculales); 74 superasterid species, representing 32 families in 11 orders, including 21 species from the basal lineages Caryophyllales, Santalales, and Ericales; and 149 superrosid species, representing 32 families in 12 orders, including four species from two basal superrosid lineages (Saxifragales and Vitales).

**Figure 1.**
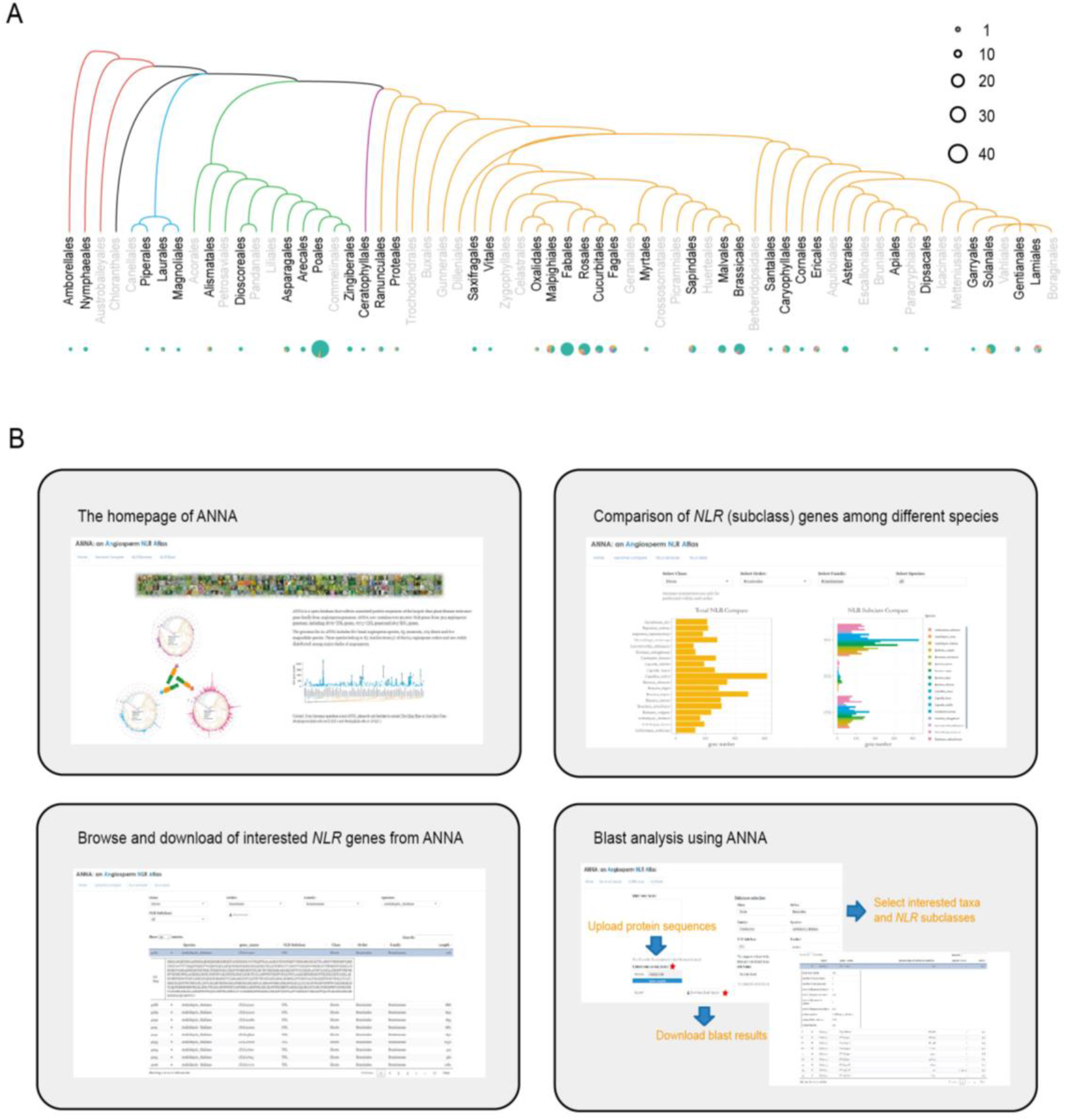
Construction of an <u>an</u>giosperm *<u>N</u>LR* <u>a</u>tlas (ANNA) covering 91,291 *NLR* genes in 304 angiosperm genomes. A) Phylogenetic relationships among angiosperm orders, constructed based on the APG IV system of flowering plant classification (Byng et al., 2016). Circle size corresponds to the number of species in each order; circle segments reflect the distribution of species among families. B) Representative screenshots of the ANNA interface, showing the major functions of ANNA: browsing, downloading, and primary comparative analysis of *NLR* datasets of interest.

We analyzed these genomes to clarify whether *TNL* gene loss occurs rarely or frequently in dicot genomes. Notably, we identified *NLR* genes in five magnoliids, a major angiosperm clade that has to date been poorly investigated due to a lack of genomic data. Exploration of these data might reveal whether the patterns of *NLR* evolution in this clade differ from those recovered in the monocots and dicots. The 91,291 *NLR* genes were unevenly distributed across the 305 genomes examined: the number of *NLR* genes per genome ranged from 5 to more than 2000, except for one genome (*Utricularia gibba*) that contained no detectable *NLR* genes. *U. gibba* represents the only known land plant genome completely lacking *NLR* genes.

All identified *NLR* genes were combined to form ANNA, a publicly available free website that allows users to browse, download, and perform primary comparative analyses of *NLR* datasets of interest (Figure 1B). In total, the ANNA dataset is comprised of 3.6 species per family; 16 families are represented by more than five genomes. ANNA thus provides a uniquely large dataset for use by the *R* gene research community, allowing users to explore various aspects of *NLR* gene evolution. The *NLR* data in ANNA were used for all subsequent analyses in this study.

### Dramatic variations in *NLR* gene numbers among angiosperms are associated with adaptions to different lifestyles

Evaluation of the 305 angiosperm genomes showed that *NLR* copy number varied tremendously among genomes, even within the same clade (Figure 2A). Ten genomes included more than 1000 *NLR* genes each (Table S2): three species in one monocot order (Poales) and seven species in five dicot orders (Gentianales, one species; Rosales, two species; Malpighiales, one species; Fagales, two species; and Myrtales, one species). This suggested several instances of lineage- or species-specific gene expansion. Consistent with this, CD-hit analysis (Huang et al., 2010) showed a high degree of sequence similarity among many of the *NLR* genes in these ten genomes (Figure S1), suggesting intense recent gene duplications. By contrast, we identified 30 genomes that possessed fewer than 50 *NLR* genes each, distributed among 21 families in 17 orders. CD-hit analysis showed that few of these genomes contained large gene clusters with high sequence similarity, indicating rare recent gene duplications (Figure S1).

**Figure 2.**
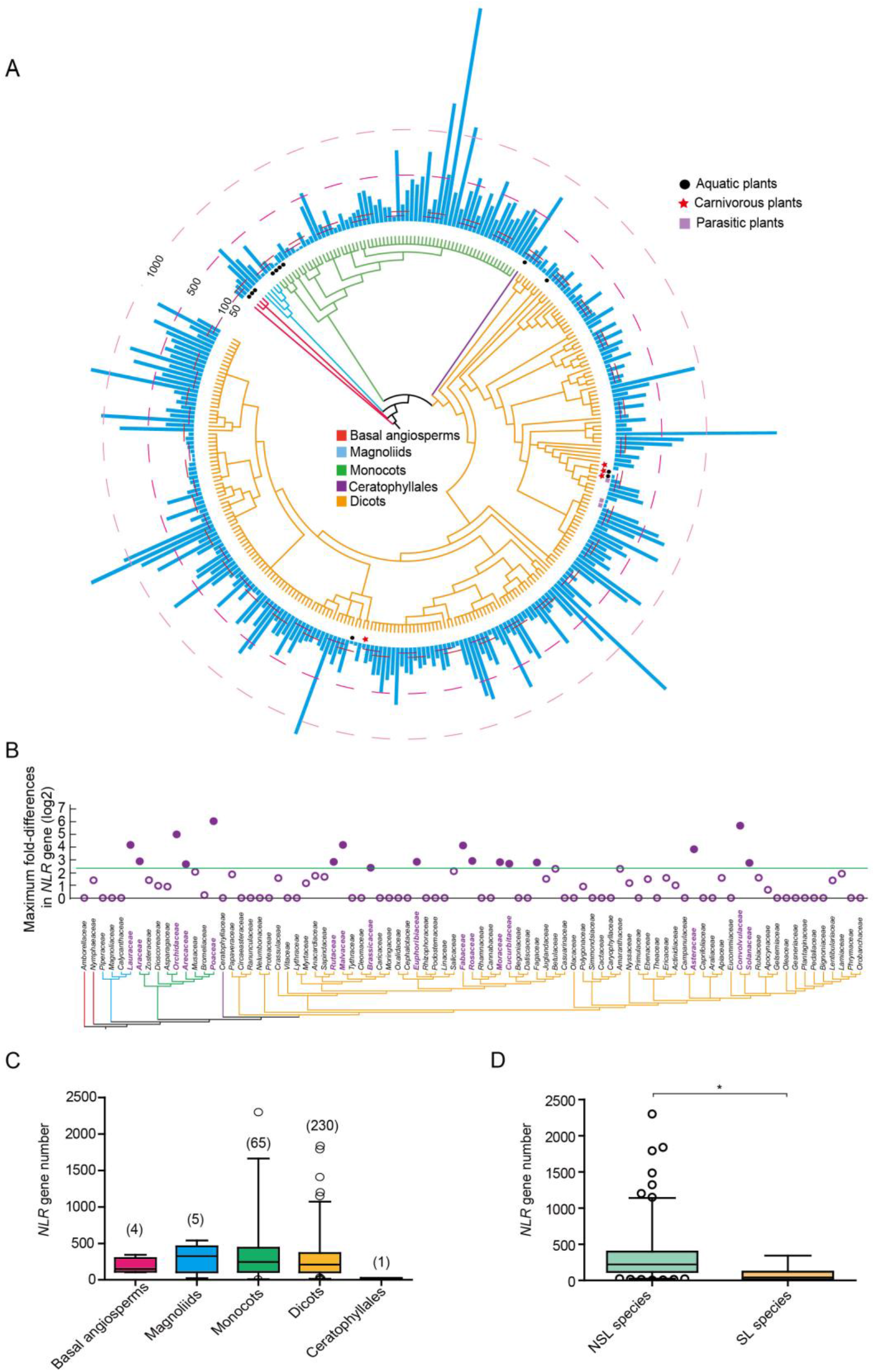
Variations in *NLR* gene number among angiosperm genomes. A) Phylogenetic relationships among the 305 angiosperm genomes included in this study, constructed following the APG IV system (Byng et al., 2016). The numbers of *NLR* genes in each genome are shown by the blue bars. B) Maximum fold-differences in *NLR* gene copy number within each angiosperm family. C) Boxplots showing *NLR* gene numbers in the genomes of the basal angiosperms (n=4), magnoliids (n=5), monocots (n=65), dicots (n=230), and Ceratophyllales (n=1). Gene numbers did not differ significantly among groups. D) Boxplots showing NLR gene numbers in the genomes of special lifestyle species (SL species, n=17) and non-special lifestyle species (NSL species, n=288). SL species includes aquatic (14 genomes), parasitic (3 genomes), and carnivorous (4 genomes, 2 of which are also aquatic) species.

The scattered distribution of species with unusually high or low numbers of *NLR* genes across the angiosperms suggested rapid *NLR* expansion and/or contraction during speciation. To test the extent to which the *NLR* gene number varied among closely related species, we calculated the fold-difference in *NLR* gene number between the species with the most *NLR* genes and the species with the least *NLR* genes within each of the 85 families. Within the 42 families for which multiple genomes were available, the fold-difference in *NLR* gene number ranged from 1.2 to more than 66 (Figure 2B). In 17 families, there was a greater than five-fold difference in interspecific *NLR* gene numbers (Figure 2B). These data suggested that *NLR* gene expansion and/or contraction can occur rapidly during speciation and may facilitate the rapid response of plants to changeable pathogenic environments. Importantly, however, there were no significant differences in *NLR* gene numbers among the five major angiosperm clades (Figure 2C), suggesting that evolutionary forces do not favor maintaining stable gene numbers in specific plant clades over evolutionary time-scales.

Across all genomes investigated, aquatic, parasitic, and carnivorous plants had significantly fewer *NLR* genes on average than other angiosperm genomes (Figure 2D). For example, of the 12 aquatic plant genomes included in this study, seven had fewer than 100 *NLR* genes, and an additional four had no more than 200 *NLR* genes (Table S2). These species included three basal angiosperms, four monocots, four dicots, and one Ceratophyllales, suggesting a convergent contraction of *NLR* genes when adapting to an aquatic lifestyle. Two aquatic dicots in the Lentibulariaceae (superasterid lineage), *Genlisea aurea* and *Utricularia gibba,* had fewer than 50 *NLR* genes, consistent with drastically shortened genomes of these species (Lan et al., 2017; Leushkin et al., 2013). However, a third Lentibulariaceae species, *U. reniformis*, which is terrestrial and has a large genome (300 Mb; (Silva et al., 2019), also had fewer than 50 *NLR* genes. Thus, the low *NLR* gene numbers in the *G. aurea* and *U. gibba* genomes might not be associated with genome size reduction and an adaptation to an aquatic lifestyle. Instead, these low numbers might be linked to a lifestyle shared by all three species: carnivory (Figure 2A). Consistent with this speculation, another carnivorous plant in the rosid lineage, *Cephalotus follicularis*, also had an unusually small number of *NLR* genes (33 genes). *NLR* copy reductions were also observed in several parasitic species (Figure 2A). The two parasitic species in the Convolvulaceae included in ANNA, *Cuscuta australis* and *C. campestris*, possessed only 15 and 16 *NLR* genes, respectively, although both have large genomes including more than 40,000 protein coding genes each (Sun et al., 2018; Vogel et al., 2018). Likewise, the genome of the parasitic dicot *Striga asiatica* contained only 89 *NLR* genes. These data indicated that convergent reductions in *NLR* genes were also associated with adaptations to a parasitic lifestyle. These examples suggested that adaptations to specific lifestyles or environments may have shaped the *NLR* profiles of the angiosperms. Additionally, the utter absence of *NLR* genes from the *U. gibba* genome shows that at least some angiosperms can survive without *NLR* genes.

### Evolutionary associations among *NLR* subclasses

Angiosperms inherited three *NLR* subclasses from their common ancestor (Shao et al., 2016). Our data revealed that the *TNL* and *CNL* subclasses together account for over 90% of all *NLR* genes in 291 out of the 305 genomes (Table S2). The remaining 14 genomes have extremely low numbers of *NLR* genes overall. *RNL* genes represented no more than 10% of all *NLR* genes in most angiosperms, reflecting the conserved signal transduction functions of these genes. *CNL* genes were the most common *NLR* genes in 268 angiosperm genomes, while *TNL* genes were the most common *NLR* genes in only 36 genomes (Figure 3A). This might be because the ancestral *CNL* lineage has gradually expanded in the angiosperms over evolutionary time, while the *TNL* lineage has remained relatively stable (Shao et al., 2016). Of the genomes in which the *TNL* genes were most common, half of them were species in the Brassicaceae. This was consistent with the ancestral expansion of the *TNL* genes in the Brassicaceae lineage (Zhang et al., 2016). The remaining 18 species in which *TNL* genes dominate were scattered among families that tend to be *CNL*-dominant, suggesting species-specific *TNL* to *CNL* ratio alterations.

**Figure 3.**
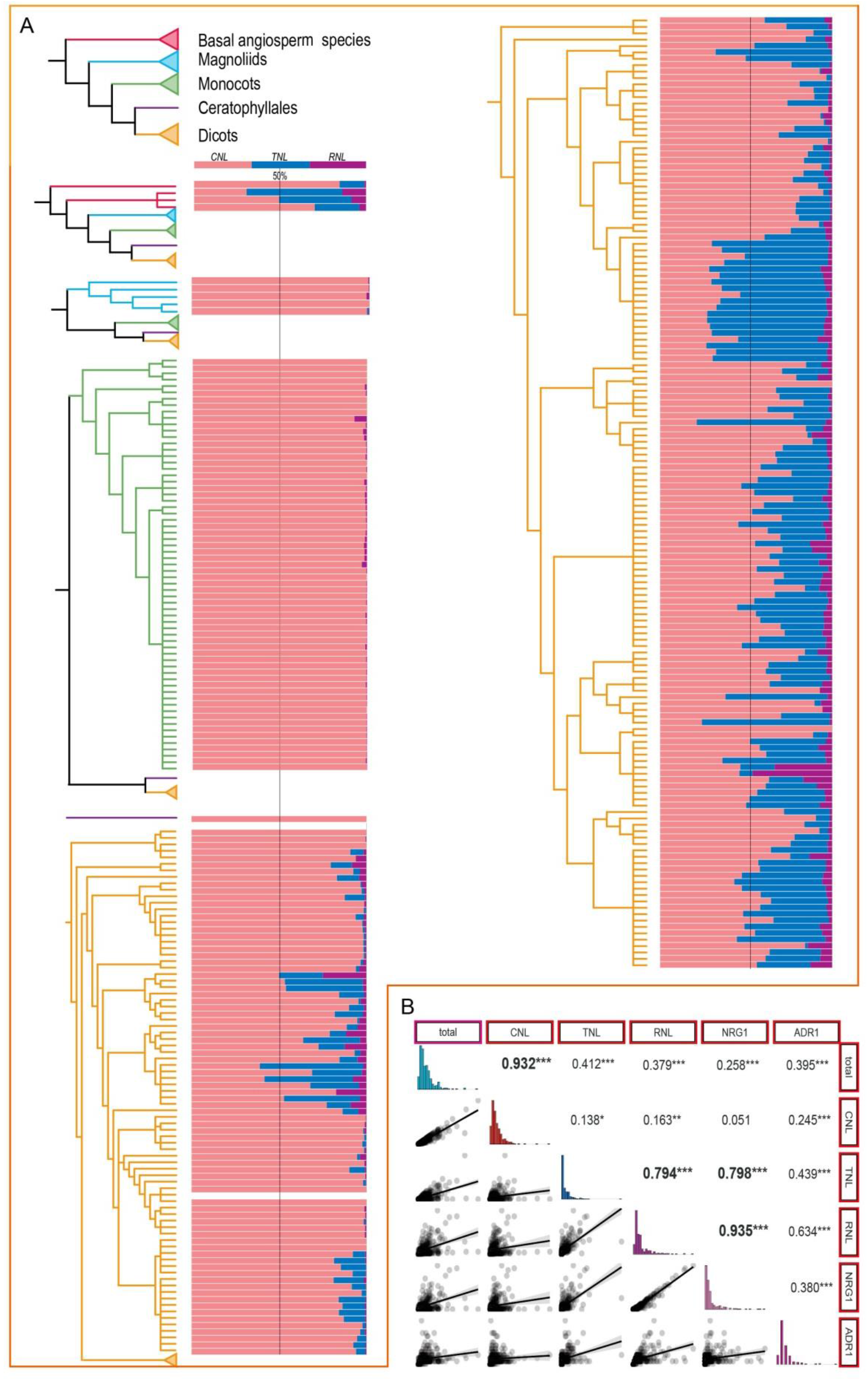
Variations in *NLR* subclass abundance among angiosperms. A) Relative proportions of the three *NLR* subclasses in the angiosperm genomes included in this study. B) Spearman correlation analysis for gene numbers among different *NLR* subclasses. The correlation coefficient is shown at top right of each graph. **, P < 0.01; ***, P < 0.001; ****, P < 0.0001.

The variations in the relative abundances of the three *NLR* subclasses among the angiosperm genomes raised two interesting questions: first, does the plant immune system require the synchronous expansion or contraction of certain *NLR* subclasses in order to respond to altered pathogenic environments; and second, have *NLR* detectors co-evolved with NLR helpers. Analysis of *NLR, CNL, TNL,* and *RNL* gene numbers identified a robust, significant correlation between total *NLR* gene number and *CNL* gene number (Figure 3B), probably due to the high relative abundance of *CNL* genes in most angiosperm genomes. The numbers of *TNL* and *RNL* genes were also significantly correlated with total *NLR* gene number, although both of these correlations were weaker than that between *NLR* gene number and *CNL* gene number. These weak correlations might be associated with the inconsistent variations in *RNL* and *TNL* gene numbers with respect to variations in *CNL* gene numbers across lineages, and the fact that neither subclass predominated in a majority of the angiosperm lineages. Consistent with this, we identified a poor correlation between *TNL* and *CNL* gene numbers, suggesting that these two *NLR* subclasses might evolve independently in response to pathogenic selection in some species. Interestingly, we observed a strong correlation between *TNL* and *RNL* copy numbers, but not between *CNL* and *RNL* copy numbers. Further dividing the *RNL* subclass into the *ADR1* and *NRG1* lineages revealed that the correlation between *RNL* and *TNL* gene copy numbers was primarily driven by the *NRG1* lineage. These results were consistent with pervious functional studies, which have shown that nearly all investigated TNL proteins are functionally dependent on RNL proteins, particularly the NRG1 lineage (Castel et al., 2019; Qi et al., 2018; Saile et al., 2020; Wu et al., 2019). In contrast, some CNL proteins could confer resistance when the *RNL* genes are deleted (Saile et al., 2020).

### Ancient and ongoing complete loss of *TNL* genes from angiosperm genomes

*TNL* loss in the common ancestor of the monocots is a familiar, long-standing hypothesis. However, genome-wide data supporting this notion have been restricted to the Poaceae and a few non-Poaceae species. Here, the data in ANNA (65 monocot genomes across nine families in six orders) provide substantial support for the hypothesis of ancestral *TNL* loss in the monocots (Figure 4A and Table S2).

**Figure 4.**
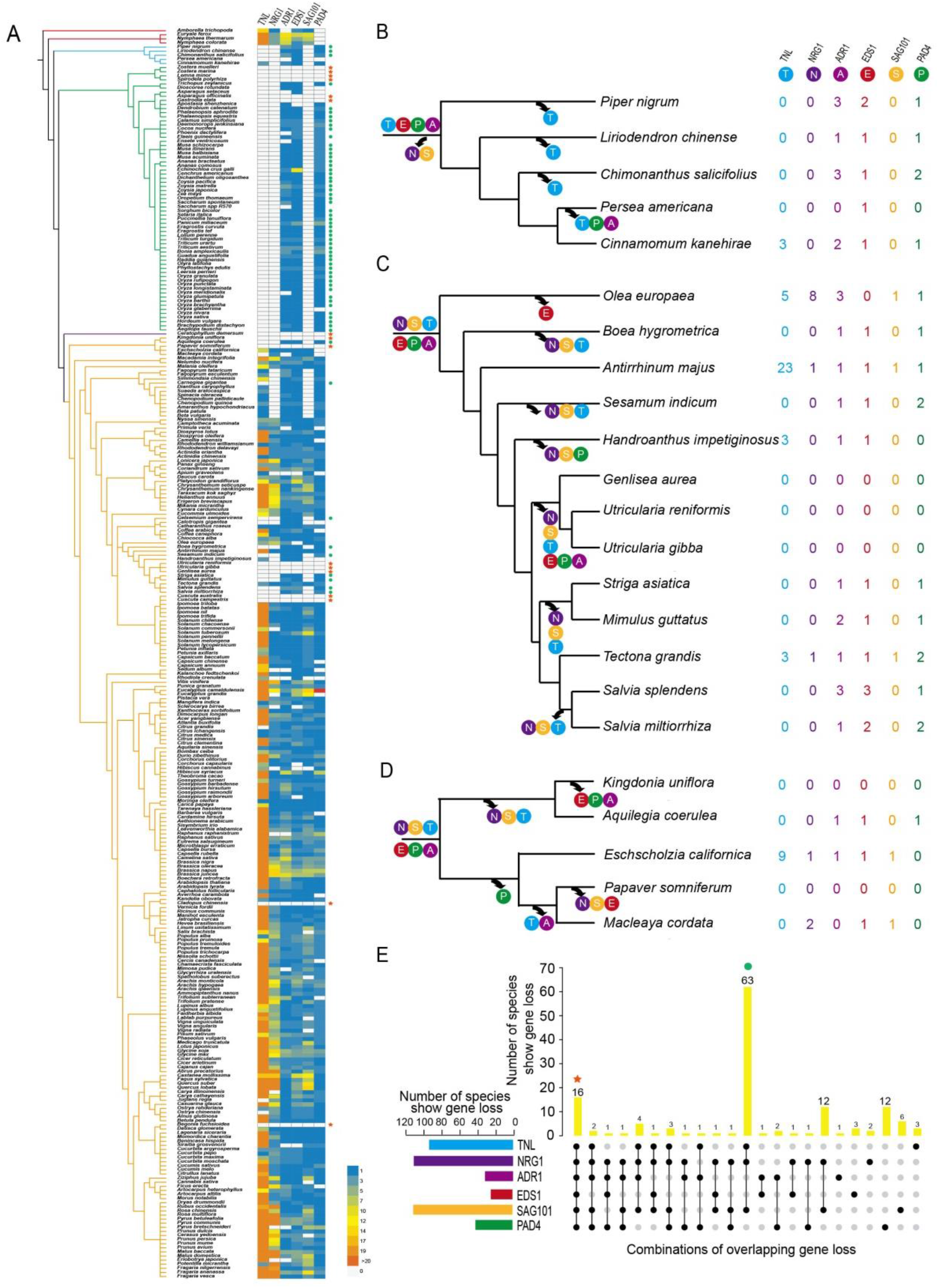
Absence of *TNL* genes and *TNL* signal transfer components in angiosperms. A) Heat map showing gene copy numbers for *TNL* and *TNL* signal transfer components (*RNL* and *EDS1* family members). Phylogenetic relationships among the 305 angiosperm genomes resolved to the order level were constructed following the APG IV system (Byng et al., 2016). Green circles indicate simultaneous losses of *TNL* and all five signal transduction components. Red stars indicate simultaneous losses of *TNL*, *NRG1,* and *SAG101*. B-D) Predicted histories of loss of *TNL* and *TNL* signal transfer components in (B) the Magnolias, (C) the Lamiales, and (D) the Ranunculales. Phylogenetic relationships among and within the three clades were constructed based on (Byng et al., 2016; Mint-Evolutionary-Genomics-Consortium, 2018; Song et al., 2017; Xue et al., 2021; Zumajo-Cardona et al., 2018). E) Numbers of species lacking *TNL* and/or various *TNL* signal transfer components. Black dots indicate gene loss, and gray dots indicate gene presence.

Recent studies have also reported the absence of *TNL* genes in a basal dicot species (*Aquilegia coerulea*) and two Lamiales species (*Sesamum indicum* and *Mimulus guttatus*), suggesting that *TNL* loss has also occurred in the dicots (Collier et al., 2011; Shao et al., 2016). *TNL* genes in angiosperms descended from only a few ancestral genes, which did not expand during long-term evolution (Shao et al., 2016). Therefore, when many genomes are surveyed, it is possible to detect additional *TNL*-lacking species among dicots that have not experienced substantial *TNL* expansion. Our data revealed that *TNL* loss indeed occurred frequently in the dicots. Of the 230 surveyed dicot species, 23 exhibited a complete loss of the *TNL* genes (Figure 4A and Table S2). These species were scattered across 18 families in nine orders and included superrosids, superasterids, and basal dicots. Moreover, of the five magnoliid genomes investigated, *TNL* genes were absent in the four more basal species, but three *TNL* genes were detected in the most derived genome, *Cinnamomum kanehirae* (Figure 4B). These results suggested that independent *TNL* losses may have occurred repeatedly throughout dicot and magnoliid diversification.

Previous studies have shown that the genomes of *S. indicum* and *M. guttatus* lack *NLR* genes, suggesting that *NLR* genes may have been lost in the common ancestor of the Lamiales (Collier et al., 2011; Shao et al., 2016). Interestingly, our data indicated an extensive but not exhaustive absence of *TNL* genes in the Lamiales order (Figure 4C). Of the 13 Lamiales species investigated, nine were lacking *TNL* genes. The remaining four species did not form a single basal clade but were instead scattered across the Lamiales phylogeny (Figure 4C). If we define a *TNL* loss event based on the presence of *TNL* genes in the sister lineage, our phylogeny suggested at least five independent *TNL* loss events throughout the diversification of the Lamiales. A similar pattern of independent *TNL* loss was also observed in the basal dicot lineage Ranunculales (Figure 4D), where three predicted *TNL* loss events resulted in the absence of *TNL* genes in four of the five genomes examined. Nine additional independent *TNL* loss events were identified: in the dicots *Carnegiea gigantea*, *Gelsemium sempervirens*, *Begonia fuchsioides*, *Cladopus chinensis*, *Juglans regia*, *Ostrya chinensis*, *Citrus grandis,* and the genus *Cuscuta,* as well as the Ceratophyllales *Ceratophyllum demersum* (Figure 4A). In total, we identified at least 23 independent *TNL* loss events across the angiosperm genomes examined, suggesting ancient, recent, and ongoing genome-wide *TNL* erasure.

### *TNL* loss is tightly associated with absence of *TNL* signal transduction components

The absence of *TNL* genes in monocots has long puzzled researchers. However, recent studies have revealed the co-absence of the *RNL-NRG1* lineage and *TNL* genes in monocots, two Lamiales species, and *A. coerulea* (Collier et al., 2011; Shao et al., 2016). In addition to *NRG1*, various other proteins play a role in TNL signal transduction, including EDS1, PAD4, SAG101, and ADR1 (Castel et al., 2019; Lapin et al., 2019; Qi et al., 2018; Saile et al., 2020; Wu et al., 2019). Across all 305 angiosperm genomes, we identified 93, 109, 29, 23, 40, and 110 lacking *TNL*, *NRG1*, *ADR1*, *EDS1*, *PAD4*, and *SAG101*, respectively (Table S2). *NRG1* and *SAG101* were the most frequently lost genes; these genes were both absent in 104 genomes. *NRG1* and *SAG101* were also the most frequently absent genes in genomes lacking *TNL* genes: of the 93 genomes lacking *TNL*, 89 had simultaneously lost either *NRG1* or *SAG101*, and 63 of these had lost both genes (Figure 4E). As expected, most of the genomes lacking *TNL* as well as *NRG1* and/or *SAG101* were monocots (65 in total). In monocots, the absence of these genes is likely the ancestral state. By contrast, in the dicots and magnoliids, the scattered losses of *TNL* as well as *NRG1* and/or *SAG101* across the phylogeny might suggest multiple independent loss events (Figure 4A). The strong correlation between *TNL* loss and the loss of the signal component(s) *NRG1* and/or *SAG101* further emphasizes the functional dependence of TNL proteins on the NRG1-SAG101 pathway.

In addition to the 89 genomes lacking *TNL* as well as *NRG1* and/or *SAG101,* more than 20 genomes carried *TNL* but lacked *NRG1* and/or *SAG101*. Conversely, only four genomes carrying *NRG1* and *SAG101* lacked *TNL* (Figure 4B). This suggested that *NRG1* and/or *SAG101* loss often preceded *TNL* loss in angiosperm genomes. This possibility was consistent with the pattern of gene loss events mapped onto species-level phylogenies for several taxa. For example, two species in the Lamiales (*Antirrhinum majus* and *Tectona grandis*) carried *TNL* as well as all five signal transduction components (Figure 4C), suggesting that the ancestral Lamiales carried all of these genes. Of the remaining 11 species analyzed in this group, nine lacked *TNL*, *NRG1,* and *SAG101*; however, one species (*Handroanthus impetiginosus*) lacked *NRG1* and *SAG101* (as well as *PAD4*), but not *TNL* (Figure 4C). This suggested that, in the Lamiales, *TNL* loss followed the loss of *NRG1* and *SAG101*. A similar pattern of gene loss was also recovered in the magnoliids (Figure 4B): our phylogeny suggested that *NRG1* and *SAG101* were absent in the ancestral magnoliid, and that the *TNL* gene was independently lost four times during subsequent diversification. One species, *Persea americana,* lost *PAD4* and *ADR1* in addition to *TNL*. Thus, in both the magnoliids and the Lamiales, our character trait mapping indicated that *TNL* was lost after *NRG1* and/or *SAG101*. Notably, although *EDS1*, *PAD4,* and *ADR1* were also occasionally lost with *TNL*, these genes were absent less frequently than *TNL, NRG1,* and *SAG101* (Figure 4E). This might be because *EDS1*, *PAD4,* and *ADR1* are required for *CNL* function and basal immunity in addition to *TNL* signal transduction, resulting in a tendency for conservation (Lapin et al., 2020; Saile et al., 2020). Overall, our results suggested that the loss of *NRG1* and/or *SAG101* might drive the erasure of *TNL* genes from angiosperm genomes.

A previous study revealed that independent small-scale deletions involving a few genes contributed to the deletion of *EDS1*, *PAD4*, and *ADR1* genes from angiosperm genomes (Baggs et al., 2020). To better understand how *SAG101* and *NRG1* were lost over evolutionary time, pairwise genomic synteny analyses were performed using both dicot and monocot species. Synteny analysis identified homologous regions surrounding *SAG101* and *NRG1* in both the monocot and the dicot species examined (Figure S2). Therefore, the repeated loss of *SAG101* and *NRG1* might be a result of small-scale or single-gene loss.

### Convergent retention of a conserved *TNL* lineage in species lacking *NRG1* and *SAG101*

Above, we reported that 27 analyzed genomes lacked *NRG1* and/or *SAG101* but retained one or more *TNL* genes (Figure 4E). Two different mechanisms may have led to the preservation of *TNL* genes in these *NRG1*/*SAG101*-deficient species. First, if the loss of *NRG1* and/or *SAG101* occurred relatively recently, *NRG1*/*SAG101*-dependent *TNL* genes may have not yet been pseudogenized or deleted from the genome. Second, some *TNL* genes may be functionally independent of the NRG1-SAG101 pathway, and therefore might be preserved despite *NRG1* and/or *SAG101* loss. To better understand how often each of these possible mechanisms are implicated in *TNL* preservation, we constructed a phylogeny of all available *TNL* genes from genomes lacking *NRG1* and/or *SAG101. TNL* sequences from *A. thaliana*, *Solanum lycopersicum,* and *Amborella trichopoda* were included to help better resolve relationships of the phylogeny (Figure 5A). If we assume that all *TNL* genes depend on the NRG1-SAG101 pathway, we would expect to observe the random loss of various *TNL* lineages in multiple angiosperm genomes after the NRG1-SAG101 pathway is destroyed. Surprisingly, one *TNL* lineage was conserved across 23 of the 27 genomes examined (Figure 5B). This *TNL* lineage also included genes from *A. thaliana*, *S. lycopersicum,* and *A. trichopoda*, suggesting that it diverged prior to angiosperm radiation. In addition, each genome within in this lineage had less than three *TNL* genes, reflecting a conserved pattern of evolution.

**Figure 5.**
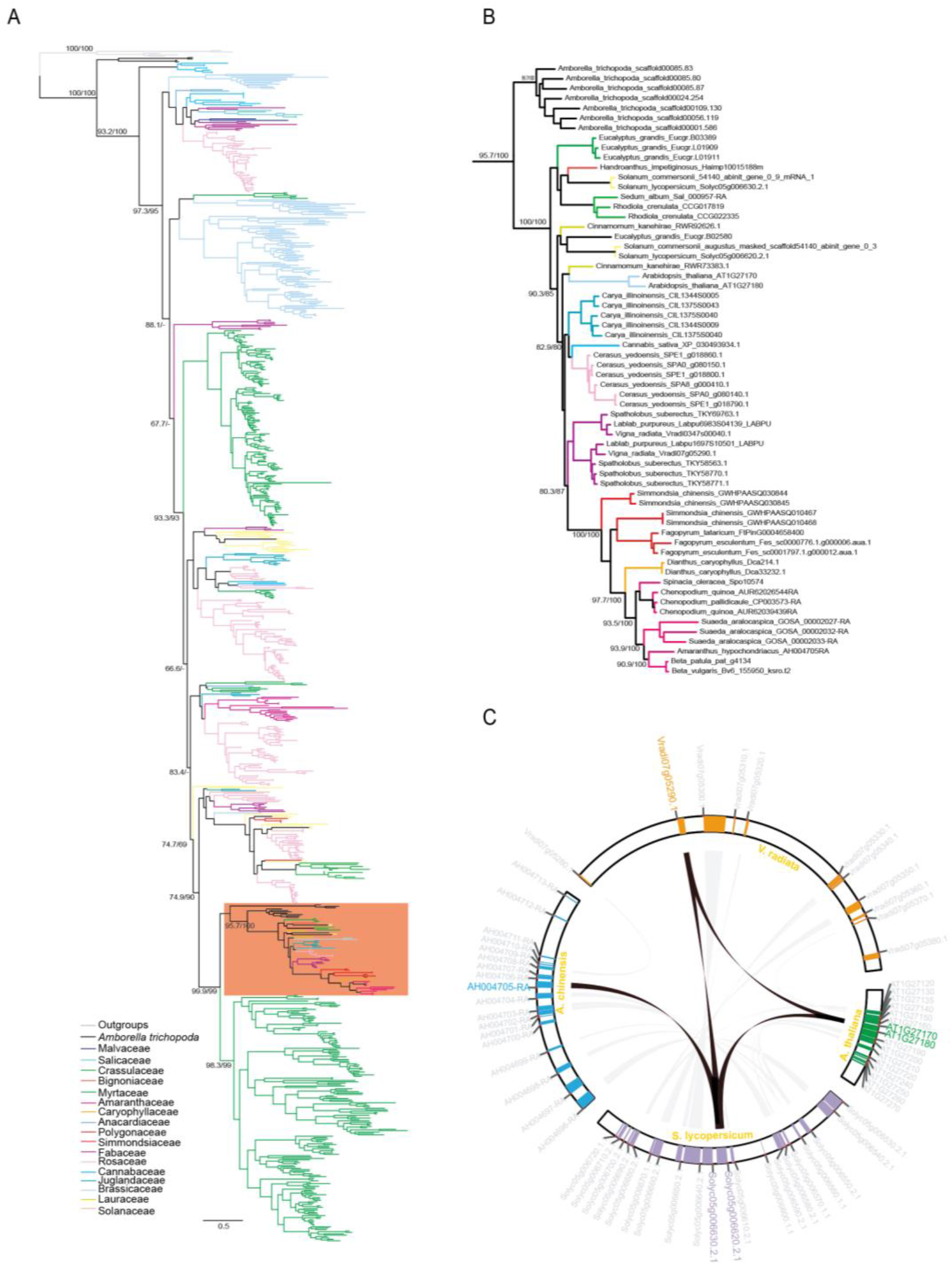
Convergent conservation of a *TNL* lineage in angiosperm species lacking the NRG1-SAG101 pathway. A) Phylogenetic analysis of *TNL* genes in 27 species carrying *TNL* but not *NRG1* and*/*or *SAG101.* The orange box indicates a conserved *TNL* lineage. Branch support values from SH-aLRT and UFBoot2 are indicated at each node. B) Expanded view of the conserved lineage highlighted in panel (A). C) Syntenic analysis of the conserved *TNL* lineage in *A*. *thaliana*, *S. lycopersicum*, *A. trichopoda,* and *V. radiata.*

To confirm that the *TNL* genes within the conserved *TNL* lineage were orthologous, we performed a syntenic analysis of the genes in this lineage from four species: the superrosid *A. thaliana*, the superrosid *Vigna radiata*, the superasterid *Actinidia chinensis*, and the superasterid *S. lycopersicum.* We found that the *TNL* genes in the conserved lineage formed well-aligned chromosomal blocks in these four species (Figure 5C), supporting an orthologous relationship among these *TNL* genes. The wide distribution of these genes, in conjunction with their highly conserved chromosomal context, suggest that this may be most conserved lineage of *TNL* genes identified in angiosperms to date. Moreover, the convergent conservation of this *TNL* lineage in species lacking *NRG1* and/or *SAG101* that diverged over 100 million years ago suggested that these *TNL* genes remain functional and may be functionally independent of the NRG1-SAG101 pathway (Figure 5B).

### Identification of candidate *TNL* signal transfer components based on evolutionary patterns

The convergent loss of *TNL* plus *NRG1* and/or *SAG101* in 89 species hinted that there might be other key signal components for *TNL* proteins, as yet unknown, that have been convergently lost. To test this hypothesis, we first identified 23 orthologous groups present in five species carrying *TNL, NRG1,* and *SAG101* (*A. thaliana*, *S. lycopersicum*, *Eschscholzia californica*, *Glycine max,* and *Olea europaea*) but absent in five species lacking all three genes (*A. coerulea*, *Musa acuminate*, *Persea americana*, *Salvia miltiorrhiza* and *Zea mays*; Figure 6A) using the online OrthoVenn2 server (Xu et al., 2019). Importantly, this analysis recovered the *NRG1*, *SAG101,* and *TNL* genes, supporting its reliability (Figure 6A, Table S5). To further investigate the loss of these 23 orthogroups in the species that had also lost *NRG1*, *SAG101,* and *TNL*, we performed a BLASTp search, querying the 20 *A. thaliana* protein sequences (representing the 20 remaining orthogroups excluding *NRG1, SAG101,* and *TNL*) against all 63 angiosperm species lacking *NRG1*, *SAG101,* and *TNL*. Five of the 20 orthogroups (clusters 4, 6, 8, 10, and 12) were also lost in over 50% of the 63 genomes lacking *NRG1*, *SAG101,* and *TNL* (Figure 6B). The GO annotations generated by the OrthoVenn2 server indicated that the *A. thaliana* genes in these five orthogroups were associated with signal transduction, transcription, the wound response, transmembrane transport activity, and the oxidative stress response (Table S5).

**Figure 6.**
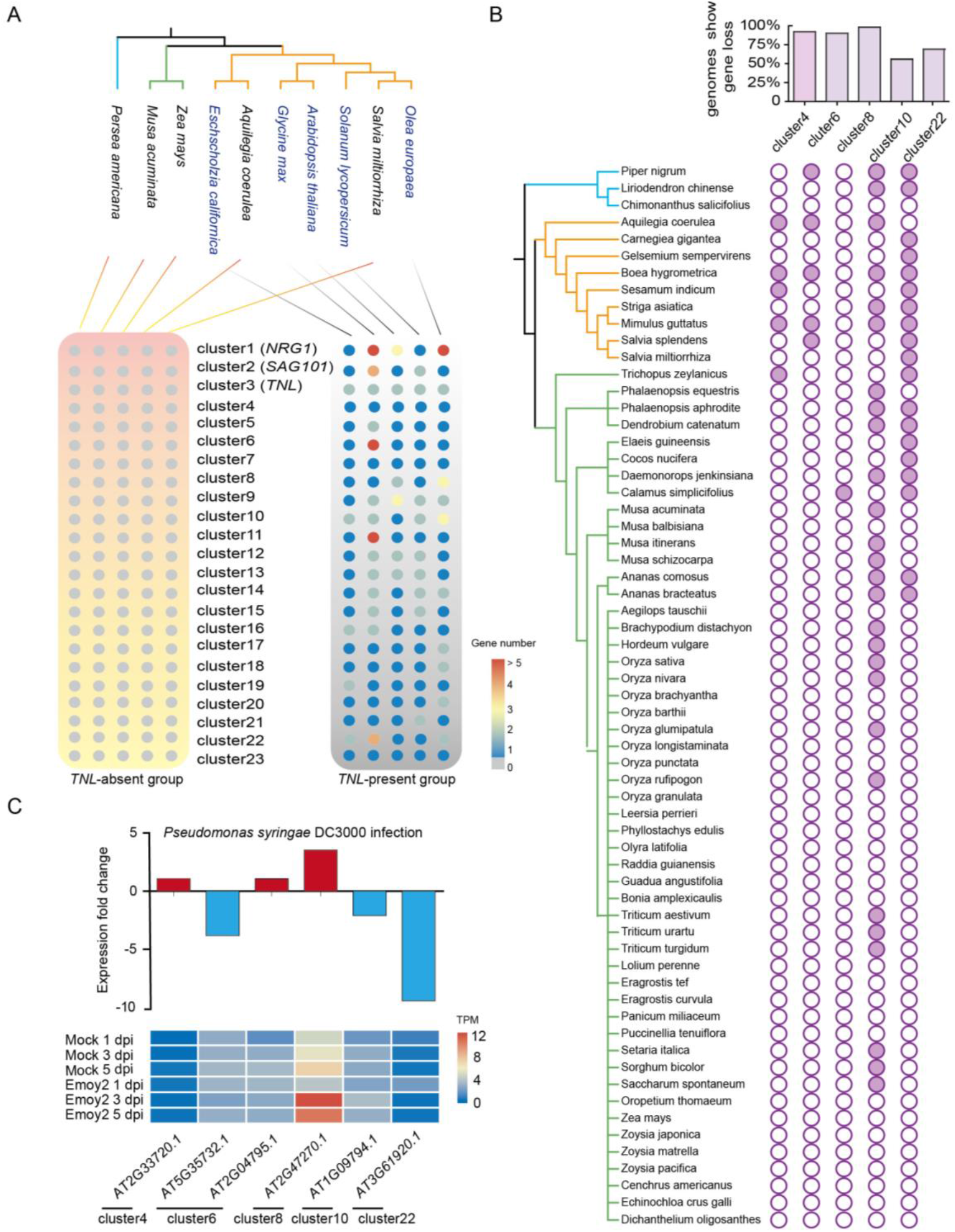
Identification of novel *TNL* signal transduction components. A) Gene clusters present in five species carrying *TNL*, *NRG1* and *SAG101* but absent in five species lacking *TNL*, *NRG1* and *SAG101*. B) Five of the 23 gene clusters were absent in more than 50% of the 63 angiosperm species lacking *TNL*, *NRG1* and *SAG101*. Empty circles indicate gene loss, while filled circles indicate gene presence. The bar graph shows the percentages of genomes lacking each gene cluster. C) The bar plot shows fold-changes in the expression levels of *A. thaliana* genes in the five clusters after *Pseudomonas syringae* DC3000 infection. Data shown are from (Asai et al., 2014a). Heat map, showing expression levels (Transcripts Per Million, TPM) of these genes after *Hyaloperonospora arabidopsidis* infection. Data shown are from (Yang et al., 2017).

To explore the potential involvement of these genes in *TNL*-triggered immunity, we performed a meta-analysis of their expression patterns in *A. thaliana* after infection with *TNL*-recognized pathogens (Asai et al., 2014b; Yang et al., 2017). We found that *A. thaliana* genes in clusters 4, 6, and 8 (AT2G33720, AT2G04795, and AT2G47270, respectively) were upregulated after infection with *Pseudomonas syringae* DC3000 (Figure 6C), which is recognized by the *TNL* protein RPS4 (Narusaka et al., 2009). AT2G47270 (cluster 4) was also upregulated when *A. thaliana* was challenged with the oomycete *Hyaloperonospora arabidopsidis* (Figure 6C), which is recognized by the TNL protein RPP4 (van der Biezen et al., 2002). AT2G47270 encodes the UPBEAT1 (UPB1) transcription factor, which harbors a bHLH domain (Li et al., 2019). A previous study showed that UPB1 regulates the expression of a set of peroxidases that modulate the balance of reactive oxygen species (ROS) (Li et al., 2019). ROS inducement is common in ETI, suggesting that UPB1 potentially acts as a signal transduction component in *TNL*-triggered immunity (Littlejohn et al., 2020). However, the exact functions of UPB1 in plant disease resistance remain to be explored.

## Discussion

Our understanding of plant *NLR* gene evolution is continuously extended as plant genomes become increasingly available (Bai et al., 2002; Meyers et al., 2003; Shao et al., 2019). Here, we developed ANNA, a database including over 90,000 *NLR* genes from more than 300 angiosperm genomes, and used this large dataset to explore the co-evolution of *NLR* subclasses, and their adaptive evolution leading to plant ecological specialization.

### *NLR* contraction tends to occur in aquatic, parasitic, and carnivorous species

Variations in *NLR* gene number among species have been widely reported. Indeed, these variations may be even more extreme than previously thought: studies of the Fabaceae, Solanaceae, Poaceae, and Brassicaceae have identified only two- to six-fold differences in *NLR* number among confamilial species, but a 20-fold difference in *NLR* number was recovered in the Orchidaceae (Luo et al., 2012; Qian et al., 2017; Shao et al., 2014; Tirnaz et al., 2020; Xue et al., 2020; Zhang et al., 2016). Here, our survey of 305 angiosperm genomes identified the greatest within-family difference in *NLR* number in the Poaceae: a 66-fold difference in *NLR* number between *T. aestivum* (2298 *NLR* genes) and *Oropetium thomaeum* (35 *NLR* genes). This pattern of extreme variation among closely related species indicates that *NLR* genes may undergo rapid and dramatic expansion and/or contraction after speciation.

It is difficult to determine the exact forces driving these species-specific *NLR* expansions and contractions because the precise qualities of the pathogenic environment during speciation are unknown. However, we identified a convergent reduction in *NLR* genes in plants with three distinct, specialized lifestyles: aquatic, parasitic, and carnivorous. That is, 11 of the 12 aquatic plants surveyed had no more than 200 *NLR* genes, and nine of these had fewer than 100 *NLR* genes. This is consistent with the findings of a previous study, which showed convergent reductions of *NLR* genes among four aquatic plants in two orders (Baggs et al., 2020). Furthermore, all carnivorous and parasitic plants surveyed exhibited convergent reductions in *NLR* gene number. These results suggested that adaptions to certain specialized lifestyles may have contributed to *NLR* reduction. Notably, no *NLR* genes were detected in the carnivorous aquatic plant *U. gibba*, demonstrating that at least some angiosperms survive without *NLR* genes.

Associations between *NLR* reduction and aquatic habitats have been observed in fern genomes (Li et al., 2018) as well as angiosperm genomes. For example, the genomes of the aquatic ferns *Azolla filiculoides* and *Salvinia cucullata* contain only one and eight *NLR* genes, respectively (Table S7). Plant-pathogen interactions are generally divided into stomatal immunity to leaf-associated pathogens and rhizospheric immunity to root-associated pathogens (Zhang et al., 2020). Although there is no empirical evidence to suggest that fewer pathogens invade aquatic plants, the anaerobic environment surrounding the roots of aquatic plants suggests that rhizospheric microbe composition might differ between terrestrial and aquatic plants. Indeed, the roots of some aquatic plants are free-floating, and thus have hardly any interaction with soil microbes. The altered or reduced interactions between roots and microbes in aquatic plants might imply that aquatic plants are less dependent on *NLR-*triggered immunity. Consistent with this, although *NLR* genes originated in the common ancestor of green plants, *NLR* genes are absent from most sequenced green algae genomes, and only began to expand after plants colonized the terrestrial environment (Shao et al., 2019). The silence of the *NLR* family in algal species during throughout several hundred million years of evolution, followed by the rapid expansion of these genes after the invasion of the terrestrial environment might reflect the reduced dependence of aquatic plants on *NLR* triggered immunity (Figure 7A).

**Figure 7.**
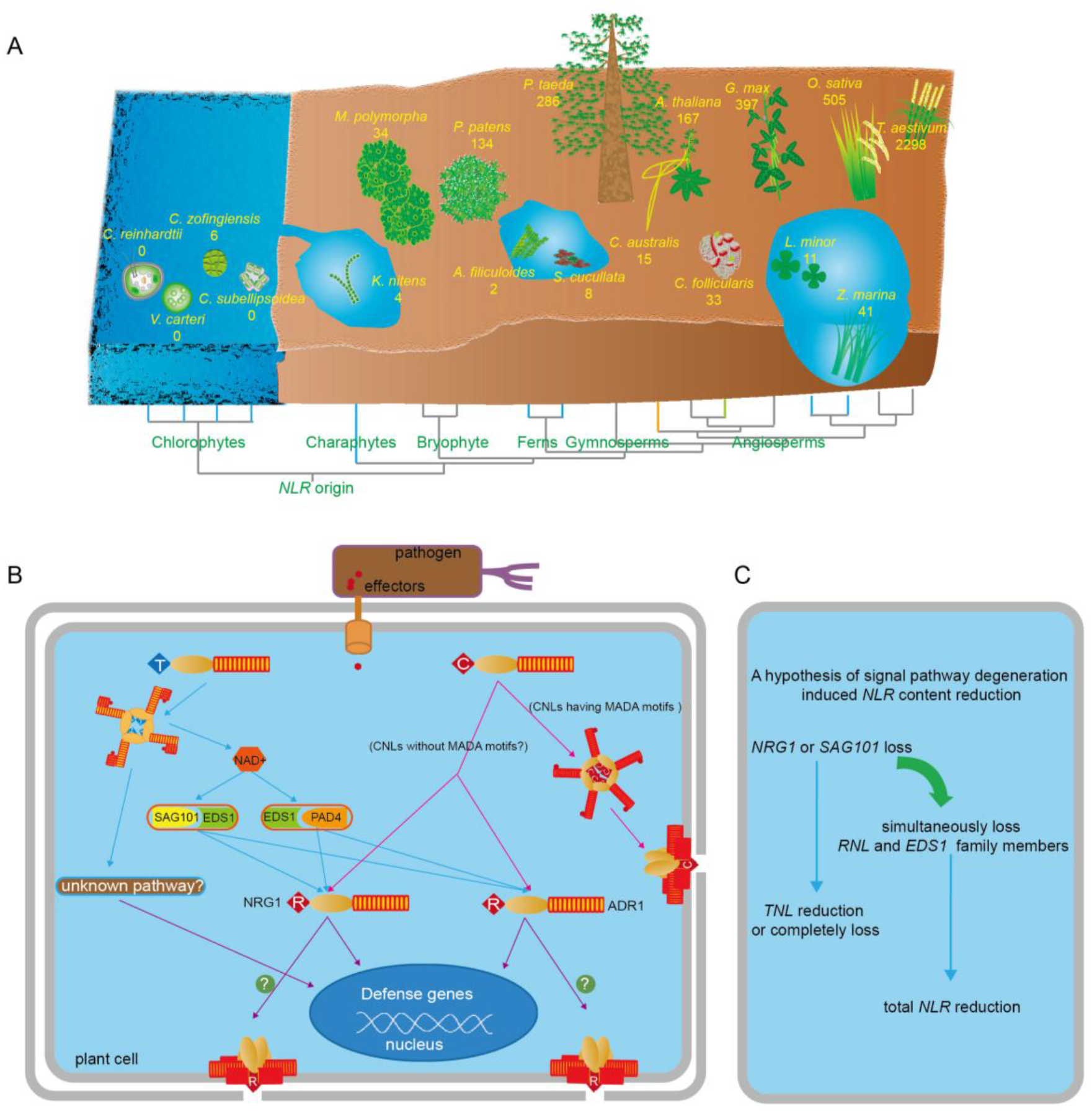
Schematic diagram showing the adaptive evolution of the *NLR* gene, leading to ecological specialization and the deletion of signal transduction components. A) A reduction in *NLR* gene number is usually coupled with adaptations to aquatic (blue branches), parasitic (yellow branch), and carnivorous (green branch) lifestyles. *NLR* gene numbers in non-vascular plants are from Shao et al (2019). B) A summary of the dependence of CNL- and TNL-triggered immunity on the EDS1 and RNL proteins. C) A hypothesized mechanism of *TNL* loss in angiosperms. EDS1 and RNL proteins are known to be important signal transduction components for most *TNL* genes. The frequent loss of *SAG101* and *NRG1* may drive *TNL* reduction and loss. However, a few *TNL* genes may be functionally independent of SAG101 and NRG1, and thus may be preserved in some species lacking a SAG101/NRG1 pathway. Additive loss of *EDS1* and *RNL* genes may cause total *NLR* number reduction.

The altered microbial environment of the roots may also explain the scarcity of *NLR* genes in parasitic and carnivorous plants. That is, parasitic plants often lack roots; some, such as *Cuscuta* spp., even lack leaves (Kaiser et al., 2015). Thus, these plants have little interaction with root (and even leaf) pathogens. Similarly, aquatic carnivorous plants are often rootless, and many hygrophytic and epiphytic carnivorous species have only weakly developed root systems (Adlassnig et al., 2005). Furthermore, most carnivorous plants live in wet soil environments that contain very limited nitrogenous material due to acidic or other unfavorable soil conditions (Xu et al., 2020). Such soil conditions may also restrict microbial growth. In addition, the leaves of most carnivorous plants form specialized trap structures that contain digestive enzymes; these enzymes may help to defend the plant from microbial invasion (Renner and Specht, 2013; Thorogood et al., 2018). Finally, carnivorous plants produce a large number of diverse secondary metabolites, many of which function as antifeedant and antifungal agents (Pavlovic and Mithofer, 2019). Thus, several characters of carnivorous plants, including their specialized root and leaf structures, low-nutrient habitat, and secondary-metabolite content, may together tend to relax the selective pressures on *NLR* genes in these taxa. In general, morphological adaptions to specialized lifestyles may have reduced the interactions between plants and microbes, leading to the contraction or elimination of *NLR* genes (Figure 7A).

### Signal transduction pathway deficiencies drive *NLR* contraction

From their common ancestor, the angiosperms inherited three *NLR* subclasses, each of which plays a unique role in PTI (Shao et al., 2016). Nearly all characterized CNL and TNL proteins participate in PTI, either by directly interacting with effector proteins, or via indirect recognition using guard or decoy mechanisms (Kourelis and van der Hoorn, 2018). This long-term arms-race drove convergent *TNL* and *CNL* expansion in the angiosperms at the Cretaceous-Paleogene (K-P) boundary (Shao et al., 2016). Interestingly, in this study, *TNL* genes were absent from the genomes of the entire monocot clade, the majority of the Magnoliids, and some of the dicots due to repeated gene loss events.

Recent studies have shown that most TNL proteins depend on RNL to confer full resistance (Castel et al., 2019; Qi et al., 2018; Saile et al., 2020; Wu et al., 2019). The dimerization or tetramerization of TNL proteins upon effector recognition activates NADase in the TIR domain, which subsequently transfers the defense signal to the downstream EDS1/SAG101 or EDS1/PAD4 complex (Ma et al., 2020; Wan et al., 2019; Zhou and Zhang, 2020). In *Arabidopsis*, the signal from EDS1 to the downstream components travels via two different pathways: one signal is passed to NRG1 to induce a hypersensitivity reaction (HR), whereas the other is passed to the ADR1 to restrict bacterial invasion (Lapin et al., 2020). Consistent with this functional analysis, our results showed that *TNL* copy number was highly correlated with *NRG1* copy number across the 305 surveyed angiosperm genomes. Furthermore, we observed the co-absence of *NRG1*/*SAG101* and *TNL* in several dozens of species. These results strongly suggested that the frequent co-absence of *NRG1*/*SAG101* and *TNL* genes is not accidental. Furthermore, the deduced prior loss of *NRG1*/*SAG101* in *TNL*-absent species suggested that the loss of *TNL* signal transfer components might be one of the driving forces underlying *TNL* loss. However, we identified a conserved *TNL* lineage in over 20 angiosperm genomes lacking *NRG1*/*SAG101*. These genomes were widely scattered throughout the angiosperm phylogeny. The convergent preservation of this anciently diverged *TNL* lineage in *NRG1/SAG101*-absent species hints that *TNL* genes might have functions independent of the NRG1-SAG101 pathway, and that these genes may trigger immunity through a different signal transduction pathway (Figure 7B). Therefore, our results suggested that *NRG1*/*SAG101* loss is the major, but not the only, factor driving *TNL* loss (Figure 7C). However, the pattern of co-absence between *TNL* and *TNL* signal pathway components helped us to identify UPB1 as a novel candidate *TNL* signal transfer component. Additional studies of this transcription factor may help to further clarify the evolution and functional mechanisms of the *TNL* genes.

While investigating the association between *TNL* loss and the loss of *TNL* signal transduction components, we identified 16 genomes (six monocots and ten dicots) lacking all five *RNL* and *EDS1* family genes (Figure 4B). It was unsurprising that species that had lost *EDS1* had also lost *TNL,* due to the critical role played by the EDS1 protein in TNL signal transduction (Lapin et al., 2020). In conjunction with the loss of *SAG101* and *NRG1,* the loss of *EDS1* would increase the likelihood of *TNL* loss. Interestingly, all 16 of these species had few *NLR* genes: 11 of these genomes had less than 50 *NLR* genes, while an additional three had no more than 100 *NLR* genes. However, *TNL* loss driven by *NRG1/SAG101* deficiency did not necessarily lead to a reduction in *NLR* gene number, as *CNL* gene expansion may occur to compensate. This pattern was observed in many monocot species, which possess many *NLR* genes but have lost *TNL* genes. Therefore, it is possible that the reduction in *NLR* number seen in the species lacking *RNL* and *EDS1* was due to a reduction in *CNL* gene number concurrent with *TNL* loss. Recent studies have shown that CNL proteins adopt at least two different mechanisms to trigger immunity (as summarized in Figure 7B). The CNL protein ZAR1 induced HR and pathogen resistance independent of both RNL lineages by assembling a polymer-protein resistosome and forming pores on the cell membrane (Saile et al., 2020; Wang et al., 2019). In addition, a conserved MADA motif was found in the CC domain of ZAR1 and other CNL proteins (Adachi et al., 2019). The motif is essential for the induction of cell death by the CNL protein in an RNL-independent manner. However, some CNL proteins do not have the MADA motif at the CC domain (Adachi et al., 2019). These CNL proteins likely do not form pores at the cell membrane, and may induce cell death by transferring the defense signal to downstream signal components. For example, the RPS2 protein, which does not harbor the MADA motif, requires ADR1 to confer full resistance (Saile et al., 2020). Several other CNL proteins are also functionally dependent on RNL (Castel et al., 2018). Therefore, the deletion of *RNL* genes from angiosperm genomes may also drive *CNL* contraction by disrupting the functions of RNL-dependent CNLs. Moreover, the convergent deletion of both *RNL* and *EDS1* family genes in certain taxa may suggest that ETI has become less important in these lineages over evolutionary time. Therefore, the reductions in *NLR* gene number observed in these species might result from the combined losses of all five component genes, as well as of other undefined immune-system components (Figure 7C).

Baggs et al. (2020) reported the convergent loss of *EDS1*, *PAD4,* and *ADR1* in four aquatic species, and hypothesized that the plant immune system co-evolved with the drought response system. Of the 16 species lacking the *RNL* and *EDS1* gene families identified in the present study, eight are indeed aquatic species: *Spirodela polyrhiza*, *Lemna minor*, *Zostera marina*, *Zostera muelleri*, *U. gibba*, *G. aurea*, *Cladopus chinensis*, and *Ceratophyllum demersum*. Although the remaining eight species (*Asparagus officinalis*, *Begonia fuchsioides*, *C. campestris*, *C. australis*, *G. elata*, *Kingdonia uniflora, Papaver somniferum*, and *U. reniformis*) are not aquatic, two are parasitic and one is carnivorous, while the remaining species have diverse lifestyles. These results emphasized that plants with aquatic and parasitic lifestyles are apt to convergently lose disease resistance signal transduction components (e.g., *RNL* and *EDS1* family genes) convergently, which may reduce the number of *NLR* genes in the genome. However, other mechanisms, yet to be characterized, apparently also result in the convergent loss of *RNL* and *EDS1* family gene as well as *NLR* contraction, as observed in the four non-aquatic, non-parasitic, and non-carnivorous species.

In conclusion, we herein established ANNA, a dataset including the *NLR* genes from more than 300 angiosperm genomes and covering most major angiosperm clades. This large dataset may serve as an essential resource for exploring various aspects of *NLR* gene evolution. Using this dataset, we revealed dramatic variations in *NLR* copy numbers among closely related species and identified notable *NLR* reductions in species adapted to aquatic, parasitic, and/or carnivorous lifestyles. The high correlation between *TNL* and *RNL* gene numbers, in addition to the high frequency of *TNL* and *NRG1* co-absence, suggested that NRG1-SAG101 pathway deficiency may drive repeated independent *TNL* losses. Our findings provide new insights into the evolution of *NLR* genes in the context of plant lifestyles and genome content variation.

## Materials and methods

### Data used in this study

The genomic sequences, annotations, and gene models for the 305 angiosperm genomes and two fern genomes were downloaded from public databases (listed in Table S1). The transcriptome data for *A. thaliana* were retrieved from previously published works (Asai et al., 2014b; Yang et al., 2017).

### Identification of *NLR* and *EDS1* family genes

*NLR* genes were identified in the 305 angiosperm genomes as described previously (Shao et al., 2015). Briefly, we first screened all protein sequences in each genome for the NBS domain (NB-ARC, Pfam: PF00931) using an HMM search, as implemented in hmmer3.0 (Johnson et al., 2010) with default parameters. The amino acid sequences of obtained *NLR* genes were then used to run a genome-wide BLASTp analysis for each genome (E-value = 1.0). All hits were further analyzed using hmmscan in hmmer3.0 against a local Pfam-A database (E-value = 10^−4^) to confirm a detectable NBS domain in each sequence. When two or more transcripts were annotated for a single gene, the longest form was selected. All obtained *NLR* candidates were classified into the *TNL*, *CNL,* and *RNL* subclasses based on a BLASTp search of their amino acid sequences against a local database containing *NLR* subclass genes representing species across the green plants (Shao et al., 2019). A similar strategy was adopted to identify EDS1 proteins. The obtained EDS1 family members were classified based on BLASTp alignments against EDS1, SAG101, and PAD4 protein sequences from *A. thaliana*, *Solanum tuberosum*, *S. lycopersicum*, *Solanum tuberosum,* and *Nicotiana benthamiana* (E-value = 1e-10).

### Sequence alignment and phylogenetic analysis

Sequence alignment and phylogenetic analysis of *TNL* genes were performed as previously described (Shao et al., 2019), with *Arabidopsis thaliana RNL* sequences as outgroups. Amino acid sequences of the NBS domain were aligned using ClustalW (Edgar, 2004) with default options (Thompson et al., 1994), and then manually corrected in MEGA 7.0 (Kumar et al., 2016). Genes with very short or divergent NBS domains were eliminated from the matrix because these interfered with fine alignment and phylogenetic analysis. Phylogenetic analyses were performed using IQ-TREE with the maximum likelihood algorithm (Nguyen et al., 2015). ModelFinder was used to estimate the best-fit model of nucleotide substitution (Kalyaanamoorthy et al., 2017). Branch support values were calculated using SH-aLRT (Anisimova et al., 2011) and UFBoot2 (Minh et al., 2013) with 1000 bootstrap replicates.

### Synteny analyses

The synteny of the conserved *TNL* lineage was analyzed in four species: *Actinidia chinensis*, *Amaranthus hypochondriacus, Arabidopsis thaliana*, and *Solanum lycopersicum*. The synteny of the *NRG1* and *SAG101* proteins was analyzed in five species: *Antirrhinum majus*, *Cladopus chinensis*, *Sesamum indicum*, *Spirodela polyrhiza*, and *Theobroma cacao*. Genome-wide pair-wise BLASTp analyses were performed between all protein sequences of the target species (E-value = 1e-10). Synteny analysis was performed with the MCScanX module of TBtools v0.6669 (Chen et al., 2020; Wang et al., 2012). Syntenic relationships were then drawn using TBtools.

### CD-hit analyses

The *NLR* protein sequences of species with more than 1000 or less than 50 *NLR* genes were analyzed using CD-HIT version 4.6 (Huang et al., 2010), with identity cutoffs of 0.8 and 0.9, respectively.

### Orthologous group identification

The online OrthoVenn2 server (https://orthovenn2.bioinfotoolkits.net, v2.0.9) (Xu et al., 2019) was used to identify orthologous gene groups in five species carrying *TNL*, *NRG1,* and *SAG101,* and in five species lacking *TNL*, *NRG1,* and *SAG101* (E-value = 1e-10). *A. thaliana* proteins from the obtained orthologous gene groups specific to species carrying *TNL*, *NRG1,* and *SAG101* were extracted to form a genomic annotation file. These sequences were used as BLASTp queries against the genomic protein sequences of all 63 species that had simultaneously lost *TNL*, *NRG1,* and *SAG101* (E-value = 1e-10).

## Acknowledgment

This work was supported by the National Natural Science Foundation of China (32070243 to Z.Q.S. and 31770245 to J.Q.C.). We thank LetPub (www.letpub.com) for its linguistic assistance and scientific consultation during the preparation of this manuscript.

## Author Contributions

Z.Q.S. and J.Q.C. conceived and designed the research. Y.L. and Z.Q.S. obtained, analyzed and inteprated the data; Z.Z., Q.L., X.M.J., J.H.T. and Y.M.Z. particpated in data analysis; Z.J. and D.C. constructed the database; Y.M.Z. and Q.W. particpated in data discussion, Y.L. and Z.Q.S. drafted the manuscript; Z.Q.S. and J.Q.C. revised the manuscript.

## Supplemental Information

**Figure S1.** Clustering of *NLR* genes from angiosperm species with A) large (> 1, 000) or B) small (< 50) number of *NLR* genes by CD-hit. The sequence identity cut-off was set as 0.8 and 0.9 respectively.

**Figure S2. Pairwise synteny analyses of five angiosperm species, showing that small-scale gene losses contribute to the independent losses of *NRG1* and *SAG101* in angiosperms.** Genes are shown as pointed colored blocks, with the point indicating gene direction, on the specified chromosome. Orthologs between two species are linked by black lines. Black arrowheads show the numbers of genes not displayed in the indicated intervals.

**Table S1.** Genomes used in this study.

**Table S2.** The gene numbers of *NLR* and *EDS1* families identified form the 305 angiosperms.

**Table S3.** Intraspecies syntenic blocks containing *NRG1* and *SAG101* genes.

**Table S4.** Intraspecies syntenic blocks containing genes from the conserved *TNL* lineage.

**Table S5.** The 23 orthogroups specific to five *TNL*-absent species identified by OrthoVenn2.

**Table S6.** Expression change of potential TNL signal transduction components in *A. thaliana* after infection by TNL-recognized pathogens.

**Table S7.** *NLR* genes identified from two ferns’ genome.

