## Supplemental Figure S1 for "An angiosperm *NLR* atlas reveals that *NLR* gene reduction is associated with ecological specialization and signal transduction component deletion"

A

B

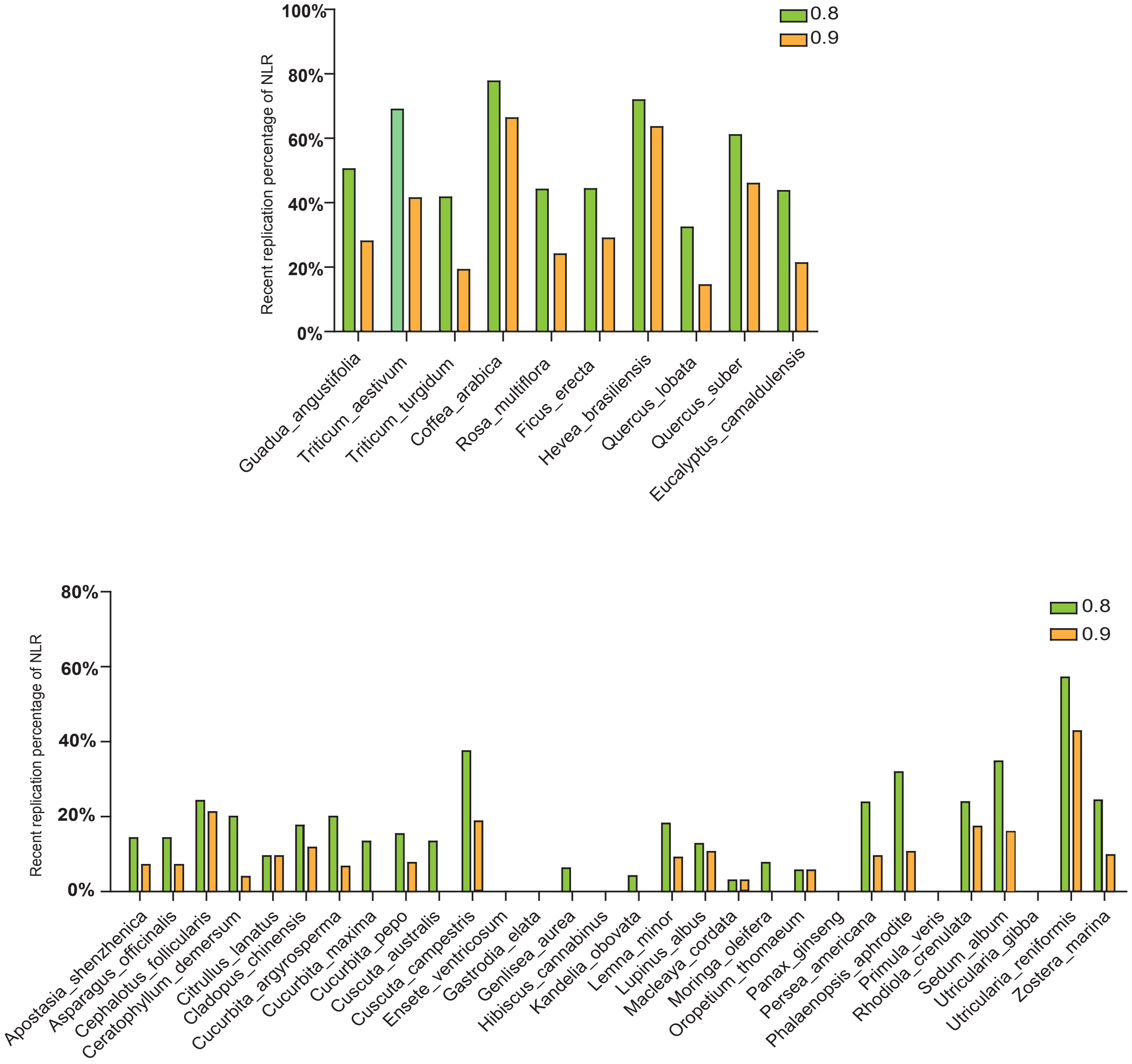


**Figure 1. Clustering of NLR genes from angiosperm species with A) large (> 1, 000) or B) small (< 50) number of NLR genes by CD-hit.** The sequence identity cut-off was set as 0.8 and 0.9 respectively.
