## Supplemental Figure S2 for "An angiosperm *NLR* atlas reveals that *NLR* gene reduction is associated with ecological specialization and signal transduction component deletion"

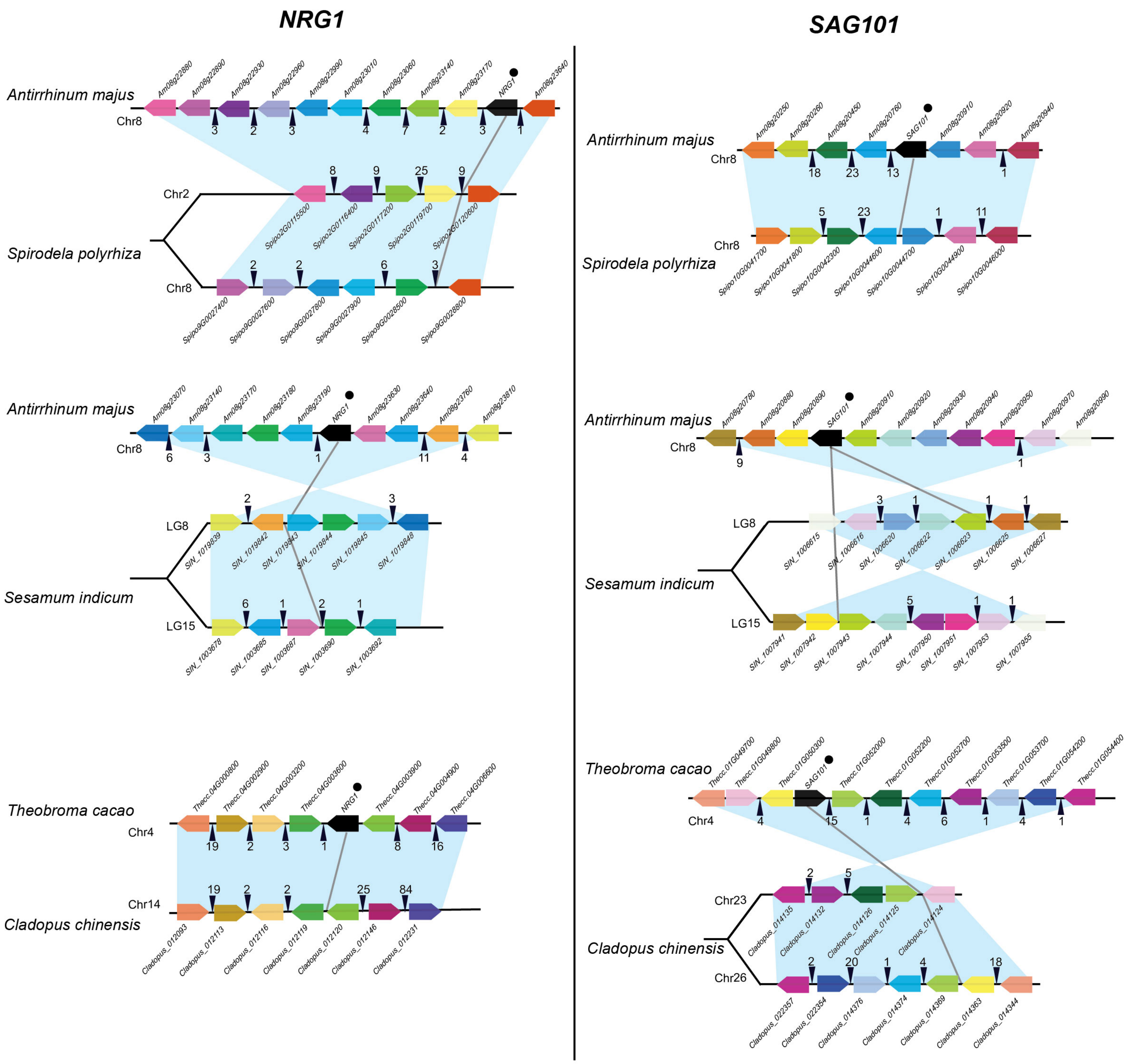


**Figure 5. Pairwise synteny analyses of five angiosperm species showing that small-scale gene losses contribute to the independent losses of *NRG1* and *SAG101* in angiosperms.** Genes are shown as pointed colored blocks, with the point indicating gene direction, on the specified chromosome. Orthologs between two species are linked by black lines. Black arrowheads show the numbers of genes not displayed in the indicated intervals.
